# Antagonistic activities of Fmn2 and ADF regulate axonal F-actin patch dynamics and the initiation of collateral branching

**DOI:** 10.1101/2020.11.16.384099

**Authors:** Tanushree Kundu, Sooraj S Das, Divya Sthanu Kumar, Lisas K Sewatkar, Aurnab Ghose

**Author notes:** Correspondence: Dr. Aurnab Ghose, Indian Institute of Science Education and Research (IISER) Pune, Dr Homi Bhaba Road, Pune 411008, INDIA.

## Abstract

Interstitial collateral branching of axons is a critical component in the development of functional neural circuits. Axon collateral branches are established through a series of cellular processes initiated by the development of a specialized, focal F-actin network in axons. The formation, maintenance and remodelling of this F-actin patch is critical for the initiation of axonal protrusions that are subsequently consolidated to form a collateral branch. However, the mechanisms regulating F-actin patch dynamics are poorly understood.

Fmn2 is a formin family member implicated in multiple neurodevelopmental disorders. We find that Fmn2 regulates the initiation of axon collateral protrusions. Fmn2 localises to the protrusion-initiating axonal F-actin patches and regulates the lifetime and size of these F-actin networks. The F-actin nucleation activity of Fmn2 is necessary for F-actin patch stability but not for initiating patch formation. We show that Fmn2 insulates the F-actin patches from disassembly by the actin-depolymerizing factor, ADF, and promotes long-lived, larger patches that are competent to initiate axonal protrusions.

The regulation of axonal branching can contribute to the neurodevelopmental pathologies associated with Fmn2 and the dynamic antagonism between Fmn2 and ADF may represent a general mechanism of formin-dependent protection of Arp2/3-initiated F-actin networks from disassembly.

## INTRODUCTION

Axonal branching is a key component in the establishment of functional neuronal circuits during development and recovery following injury. The process allows a single neuron to make multiple synaptic contacts and is critical for the development of accurate connectivity, structural plasticity and state-dependent recruitment of neuronal groups (Armijo-Weingart and Gallo, 2017; Gibson and Ma, 2011; Kalil and Dent, 2014; Low and Cheng, 2006; Rockland, 2018).

A major mechanism of axonal branching involves the development of interstitial collateral branches. The process of generating axon collaterals is a multistep process involving both extrinsic and intrinsic cues (Armijo-Weingart and Gallo, 2017; Chalmers et al., 2016; Katz, 1985; Menon and Gupton, 2018). Collaterals which appear *de novo* from the axon shaft can be induced by extracellular cues, such as netrin1 (Dent et al., 2004; Lebrand et al., 2004) nerve growth factor (NGF) (Gallo and Letourneau, 1998), brain-derived growth factor (BDNF) (Danzer et al., 2002; Marler et al., 2008). On the other hand, factors such as semaphorin 3A (Dent et al., 2004) and slit1a (Campbell et al., 2007) act locally to suppress branch formation. These cues trigger downstream signal transduction pathways, which impinge on the underlying cytoskeletal dynamics to form a branch.

The process of axonal branching begins with the formation of a focal F-actin patch (Andersen et al., 2011; Gallo, 2006; Ketschek and Gallo, 2010; Orlova et al., 2007). These transient patches are a spontaneous accumulation of branched actin mainly regulated by the WAVE1 activated Arp2/3 complex that nucleates branched actin filaments (Spillane et al., 2012). Only a small proportion of these patches forms an axonal protrusion (Armijo-Weingart and Gallo, 2017; Ketschek and Gallo, 2010; Orlova et al., 2007). The proposed model for the conversion of a patch into a protrusion involves the convergence of branched actin filaments which are rapidly extended by the activities actin elongators such as Ena/VASP proteins and formins (Armijo-Weingart and Gallo, 2017; Yang and Svitkina, 2011). Other activities involve bundling of extending F-actin filaments, membrane remodelling and finally consolidation by microtubule invasion. While the Ena/VASP family of proteins has been demonstrated to promote collateral branching (Dwivedy et al., 2007), not much is known about the function of formins in the generation of axonal protrusions to initiate collateral branching (Kawabata Galbraith and Kengaku, 2019).

Fmn2 is expressed in the developing and the mature nervous systems across vertebrate phyla (Dutta and Maiti, 2015; Leader and Leder, 2000; Nagar et al., 2020; Sahasrabudhe et al., 2016). While Fmn2 has been implicated in neurodevelopmental disorders, intellectual disability, age-related dementia, microcephaly and corpus callosum agenesis in humans (Agis-Balboa et al., 2017; Almuqbil et al., 2013; Anazi et al., 2017; Gorukmez et al., 2020; Law et al., 2014; Marco et al., 2018; Perrone et al., 2012), its function in neurons is only recently being discovered. At the neuronal growth cone, Fmn2 regulates filopodial dynamics and stability of adhesion complexes to mediate axonal outgrowth (Ghate et al., 2020; Sahasrabudhe et al., 2016). Fmn2 also regulates microtubule dynamics and influences growth cone turning (Kundu et al., 2020). Further, Fmn2 may influence dendritic spine density in hippocampal neurons (Law et al., 2014).

In this study, we identify a novel role for Fmn2 in initiating axon collateral branches. We show that Fmn2 is critical for the maintenance and stability of that F-actin patches that generate axonal protrusions. This specialised axonal F-actin network is protected from disassembly by actin depolymerising factor (ADF) by Fmn2-mediated F-actin assembly.

## RESULTS

### Fmn2 promotes initiation of axon collateral branching in spinal neurons

Formins are involved in neuronal growth and axon guidance during development (Kawabata Galbraith and Kengaku, 2019). Several formin family members localise to growth cone filopodia and regulate filopodial dynamics (Barzik et al., 2014; Rama et al., 2018; Szikora et al., 2017). The F-actin nucleation/elongation function of formins are also implicated in the dynamics of axonal actin ‘hotspots’ (Ganguly et al., 2015). Fmn2 function in growth cones is necessary for axonal outgrowth and growth cone filopodial stability (Ghate et al., 2020; Sahasrabudhe et al., 2016). As a majority of axon collateral branches arise from a filopodia-like protrusion from the axonal shaft, we investigated if the formin, Fmn2, was involved on axon collateral branching.

To evaluate the role of Fmn2 in axonal branching, Fmn2 was depleted in chick spinal neurons using well-characterised, specific translation-blocking morpholinos targeting Fmn2 (Fmn2-MO; see Materials and Methods). Phalloidin staining of F-actin (Figure 1 A,B) revealed a drastic reduction of collateral axonal protrusions (Figure 1 C) in neurons transfected with Fmn2-MO (0.118 ± 0.016 branches per µm) compared to control morpholino (Ctl-MO) transfected neurons (0.193 ± 0.02 branches per µm).

**Figure 1.**
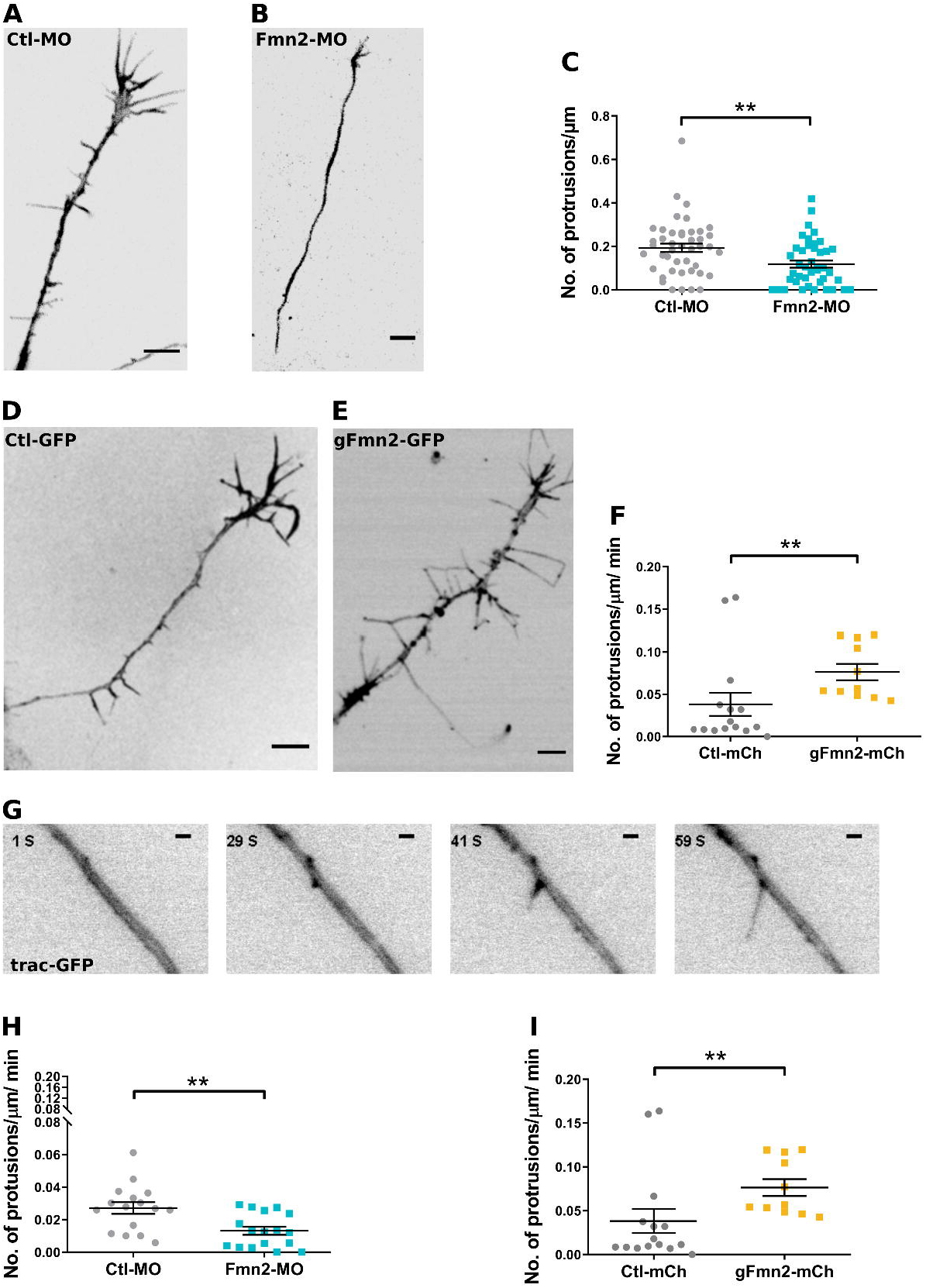
Fmn2 promotes collateral protrusions in axons by increasing rate of branch formation. Representative images of neuron with collateral protrusions after co-transfection of either Ctl-MO (A) or Fmn2-MO (B) and pCAG-GFP. The neurons were fixed and stained with phalloidin after 24 hours of incubation. (C) Graphical representation of the number of protrusions (Ctl-MO, n = 44 and Fmn2-MO, n = 42; p=0.003). (D) and (E) show representative phalloidin-stained micrographs of collateral protrusions in GFP and gFmn2-GFP overexpressing neurons 24 hours post-transfection. (F) Graphical representation of the number of protrusions (GFP, n = 36 and gFmn2-GFP n = 33; p<0.0001). (G) Four frames from a time-lapse recording of the development of a collateral protrusion from a F-actin patch. Tractin-GFP (trac-GFP) was used to label F-actin. (H) Quantification of the frequency of new protrusion formation in in morpholino treated neurons (Ctl-MO, n= 16 axons; Fmn2-MO, n= 17 axons; p=0.004). (I) Frequency of protrusion initiation in Fmn2/GFP overexpressing neurons (Ctl-GFP, n= 15 axons; Fmn2-GFP, n= 11 axons; p=0.004). The data are obtained from at least three independent experiments. All the values were plotted with the mean +/-SEM indicated. Data was analyzed using the two-tailed Mann-Whitney U test. * *, p<0.01. Scale bar:10 µm (A,B,D,E) and 1 µm (G).

Conversely, the overexpression of GFP-tagged chick Fmn2 (gFmn2-GFP; Figure 1 E) doubled the frequency of collateral branching in axons (0.246 ± 0.022 branches per µm; Figure 1 F) as compared to control GFP-expressing neurons (0.128 ± 0.015 branches per µm; Figure 1 D).

To observe axonal collateral formation, live-imaging was undertaken with neurons transfected with the GFP-tagged F-actin probe, F-tractin (trac-GFP; Figure 1 G). As reported earlier (Andersen et al., 2011; Ketschek and Gallo, 2010; Orlova et al., 2007), axon collateral branching was preceded with the emergence and elaboration of a juxta-membrane F-actin patch. The F-actin patch, extended into the membrane protrusion to initiate collateral branching. Time-lapse tracking of F-tractin in Fmn2 depleted axons revealed a significantly reduced frequency of branch initiation (0.013 ± 0.003 protrusions/µm/ min; Figure 1 H) compared to Ctl-MO co-transfected axons (0.027 ± 0.004 protrusions/µm/ min; Figure 1 H). Conversely, F-tractin-tagged axons overexpressing mCherry-tagged chick Fmn2 (gFmn2-mCh) exhibited a significantly higher rate of branch initiation (0.076 ± 0.009 protrusions/µm/ min; Figure 1 I) than axons overexpressing mCherry (0.038 ± 0.013 protrusions/µm/ min; Figure 1 I). The live imaging experiments allow us to identify Fmn2 function in protrusion initiation rather than other function, like stabilizing existing protrusions.

Thus, Fmn2 protein expression dynamically regulates the initiation of axon collateral branching in spinal neurons and highlights a novel developmental function for Fmn2 in neuronal development.

### Fmn2 localises to the base of axonal protrusions and co-localises with actin patches

Co-expression of tagged gFmn2 and F-tractin revealed the accumulation of Fmn2 in chevron-shaped structures at the axonal membrane base of axonal protrusions (Figure 2 A). Time-lapse imaging of the axon shaft revealed the co-localisation of Fmn2 to the developing axonal F-actin patch that could subsequently contribute to the generation of an axonal protrusion. Following the development of the protrusion, Fmn2 remained accumulated at the base of the protrusion for several minutes after the branch had been initiated (Figure 2 B).

**Figure 2.**
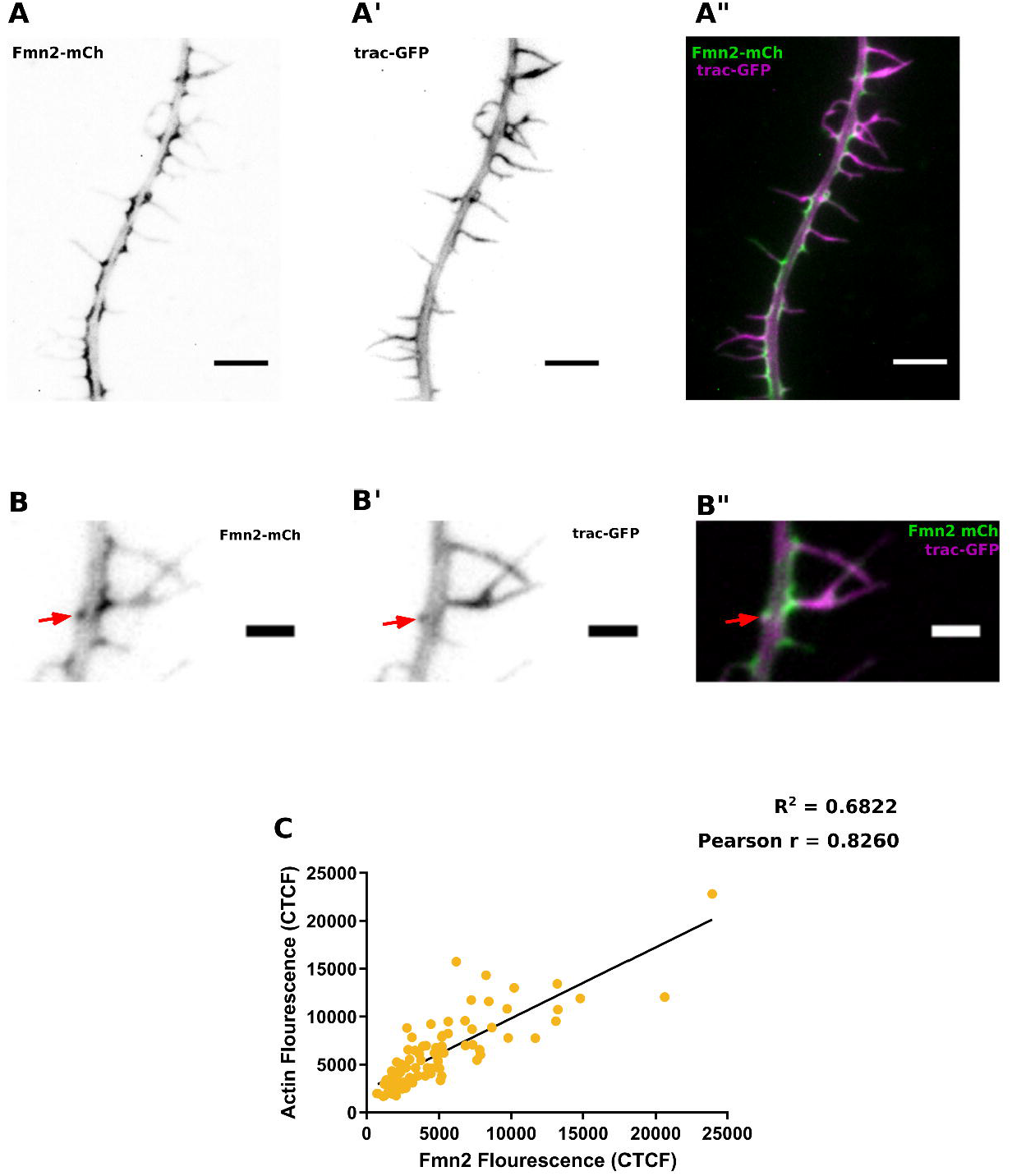
Fmn2 accumulates at the base of collaterals and co-localises with axonal F-actin patches. (A-A”) Representative micrographs of Fmn2 (grayscale in A and green in merge) and actin (grayscale in A’ and magenta in merge) and the overlay of the two channels in A”. Fmn2 forms chevron shaped structure at the base of every axonal protrusion. (B-B”) Representative micrographs from dual-colour time-lapse imaging showing Fmn2 (grayscale in B and green in merge) and actin (grayscale in B’ and magenta in merge) and the overlay of the two channels in B”. Red arrow indicates the F-actin with Fmn2, which eventually initiates a protrusion. Fmn2 was found to colocalise with F-actin patches and the co-localisation is quantified in (C). The values are plotted as XY data points with a linear regression slope (n = 86; Pearson’s correlation coefficient = 0.8260; R^2^ = 0.6822; p<0.0001). The data are from three biological replicates. CTCF, corrected total cell fluorescence. Scale bar: 2 µm (A) and 5 µm (B).

Analysis of F-tractin labelled patches revealed a significant correlation of Fmn2 co-localisation to axonal F-actin patches (Figure 2 C; Pearson correlation 0.82; R*2*= 0.68). These results revealed localisation of Fmn2 to axonal F-actin patches that may give rise to axonal branches and the accumulation of Fmn2 at the base of axonal protrusions.

### Axonal F-actin patch dynamics, but not formation, is regulated by Fmn2

Axonal F-actin networks are expected to exist in a dynamic equilibrium between nucleation activities and depolymerising activities. A proportion of axonal F-actin patches induce the development of axon collateral branches by initiating axonal protrusions. However, little is known about the dynamics of these axonal F-actin networks. To identify Fmn2 function in axonal F-actin patches, we dynamically visualized these networks used fluorophore-tagged F-tractin as a probe.

Neither Fmn2 depletion nor overexpression (both conditions that alter the extent of collateral branching) changed the density of F-actin patches in axons (Figure 3 A,B). While the above data suggested the lack of involvement of Fmn2 in initiating the formation of F-actin patches, the analysis of patch sizes revealed a positive correlation with the expression levels of Fmn2. The size of the F-actin patches was significantly reduced upon Fmn2 depletion (0.152 ± 0.012 μm^2^; Figure 3 C) compared to Ctl-Mo treated neurons (0.181 ± 0.012 μm^2^; Figure 3 C). On the other hand, compared to controls (0.135 ± 0.015 μm^2^; Figure 3 D), overexpression of Fmn2 led to larger patch sizes (0.178 ± 0.011 μm^2^; Figure 3 D).

**Figure 3.**
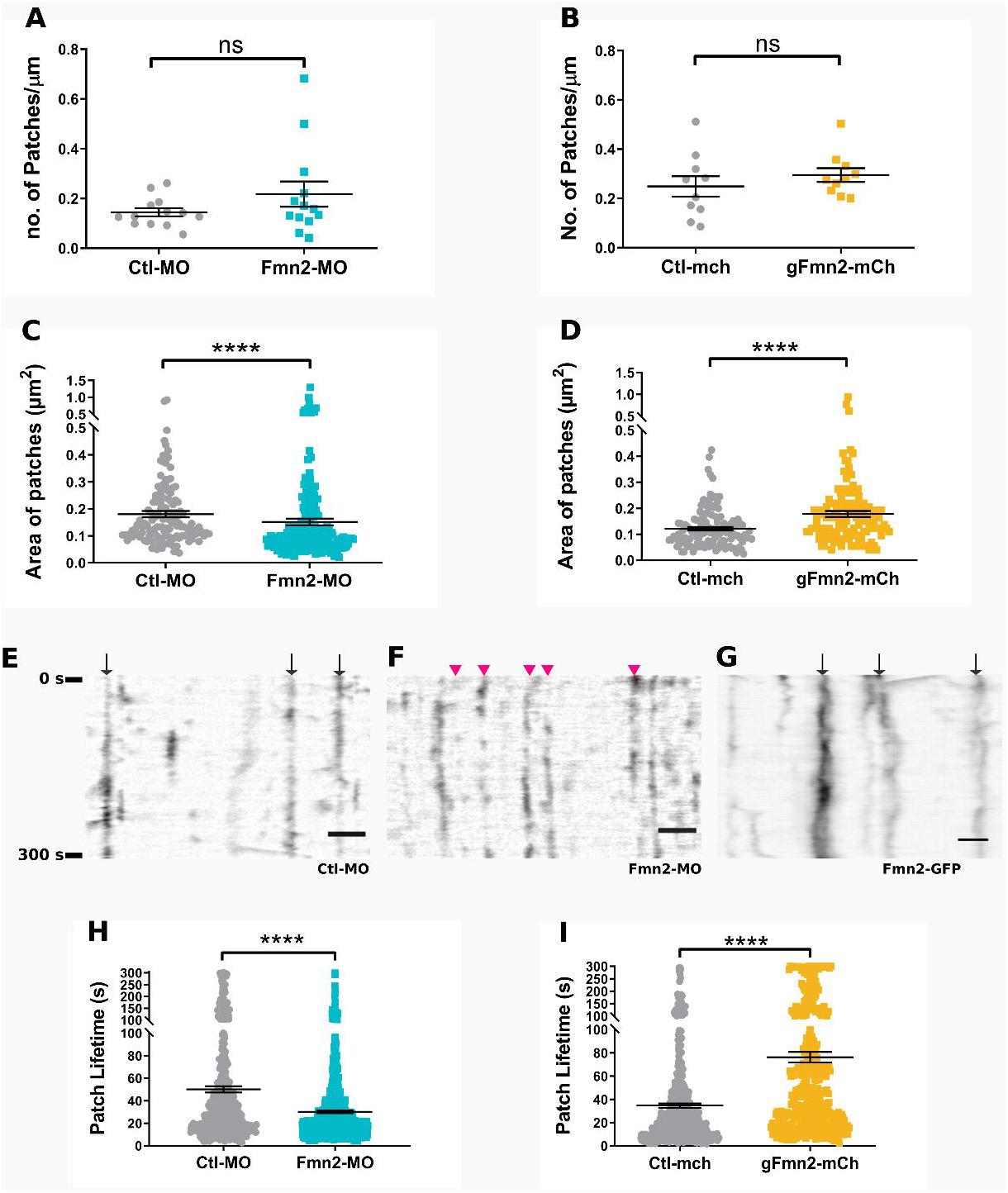
Fmn2 is necessary for patch stability but does not affect actin patch formation. (A) Density of F-actin patches in the entire axon are unchanged between Ctl-MO (n = 13 axons) and Fmn2-MO (n = 13 axons; p=0.3897) transfected neurons. (B) Overexpression of mCherry-tagged chick Fmn2 (gFmn2-mCh; n = 10) does not change the density of axonal F-actin patches compared to fluorophore (Ctl-mCh) expression neurons (n= 10 axons; p=0.3930). (C) The area of axonal F-actin patches is lower in Fmn2-MO transfected neurons (n = 189) compared to controls (n = 128; p<0.0001). Conversely, Fmn2 overexpression (gFmn2-mCh; n = 125; p<0.0001) increases F-actin patch size relative to control neurons (Ctl-mCh; n = 121). (E-G) Representative kymographs, from Ctl-MO (E), Fmn2-MO (F) and Fmn2 (G) transfected neurons indicating F-actin patches lifetimes. The pink arrow heads in (F) indicate the repeated appearance and disappearance ‘blinking’ of the F-actin patch in the same location over time. The quantification of actin patch lifetimes is shown in (H) for Ctl-MO (n = 490) and Fmn2-MO (n = 693; p<0.0001), and in (J) for Ctl-mCh (n = 576) and Fmn2-mCh (n = 379; p<0.0001). The data are obtained from three independent experiments. All the values were plotted with the mean +/-SEM indicated. The two-tailed Mann-Whitney U test was used as a test of significance. ns, p>0.05; * * * *, p<0.0001. Scale bar: 2 µm.

We analysed kymographs generated from the time-lapse images of tractin-GFP to assess the dynamics of the F-actin patches (Figure 3 E-G). Reduced levels of Fmn2 drastically shortened the lifetime of the F-actin patches from 53.6 0 ± 4.821 s in controls to 33.93 ± 2.136 s in Fmn2-MO transfected neurons (Figure 3 F,H). The analysis also revealed that Fmn2 knockdown resulted in multiple cycles of patch formation and disappearance occurring in the same region of the axon (Figure 3 F, red arrowheads), leading to a ‘blinking’ phenomenon. This observation underscores the dynamic balance of F-actin patch assembly/disassembly and implicates Fmn2 in patch maintenance. The fact that the patch ‘blinks’ at the same location suggests the presence of a nucleating core that is then elaborated by assembly promoting factors including Fmn2. Upon depletion of Fmn2, the balance shifts in favour of disassembly. Repeated nucleation is also suggestive of the lack of involvement of Fmn2 in F-actin patch initiation.

Overexpression of gFmn2 increased the F-actin patch lifetime (76.28 ± 4.477 s; Figure 3 G,H) compared to controls (34.65 ± 1.895 s; Figure 3 G,H), suggesting a stabilising function.

In order to relate patch sizes and lifetimes to the initiation of axonal collaterals, we compared these parameters between F-actin patches that result in axonal protrusion and to all patches. F-actin patches that subsequently resulted in an axonal protrusion were relatively larger and were long-lasting (Figure S1 A-D). In fact, the few patches that resulted in filopodia, despite Fmn2 knockdown, were comparable in size and lifetime with those initiating protrusions in controls (Figure S1 A-D).

Collectively, our data indicate the regulation of F-actin patch dynamics by Fmn2. F-actin patches need to elaborate to a competent size and lifetime before it can initiate an axonal protrusion. While not inducing patch formation, Fmn2 appears to be essential in the maintenance of F-actin patches allowing enough patches to reach protrusion initiation competence.

### F-actin nucleation activity of Fmn2 is required for patch stability and branch initiation

*In vitro* studies have shown that Fmn2 nucleates F-actin filaments and facilitates elongation by barbed-end actin polymerization (Montaville et al., 2014; Montaville et al., 2016). Data from the fly orthologue of Fmn2, *Cappuccino*, indicated that the mutation of a single isoleucine residue to alanine abrogated its F-actin nucleation activity (Quinlan et al., 2007; Roth-Johnson et al., 2014). Sequence alignment revealed that this residue was conserved across taxa (Figure 4 A,B). We mutated this conserved isoleucine at position 1226 to alanine in mouse Fmn2 (mFmn2-IA). Mouse Fmn2 (mFmn2-FL) cDNA does not have the binding site for the translation-blocking morpholino’s against chick Fmn2, thus mFmn2-FL and mFmn2-IA were co-expressed along with Fmn2-MO in rescue experiments to evaluate Fmn2 function.

**Figure 4.**
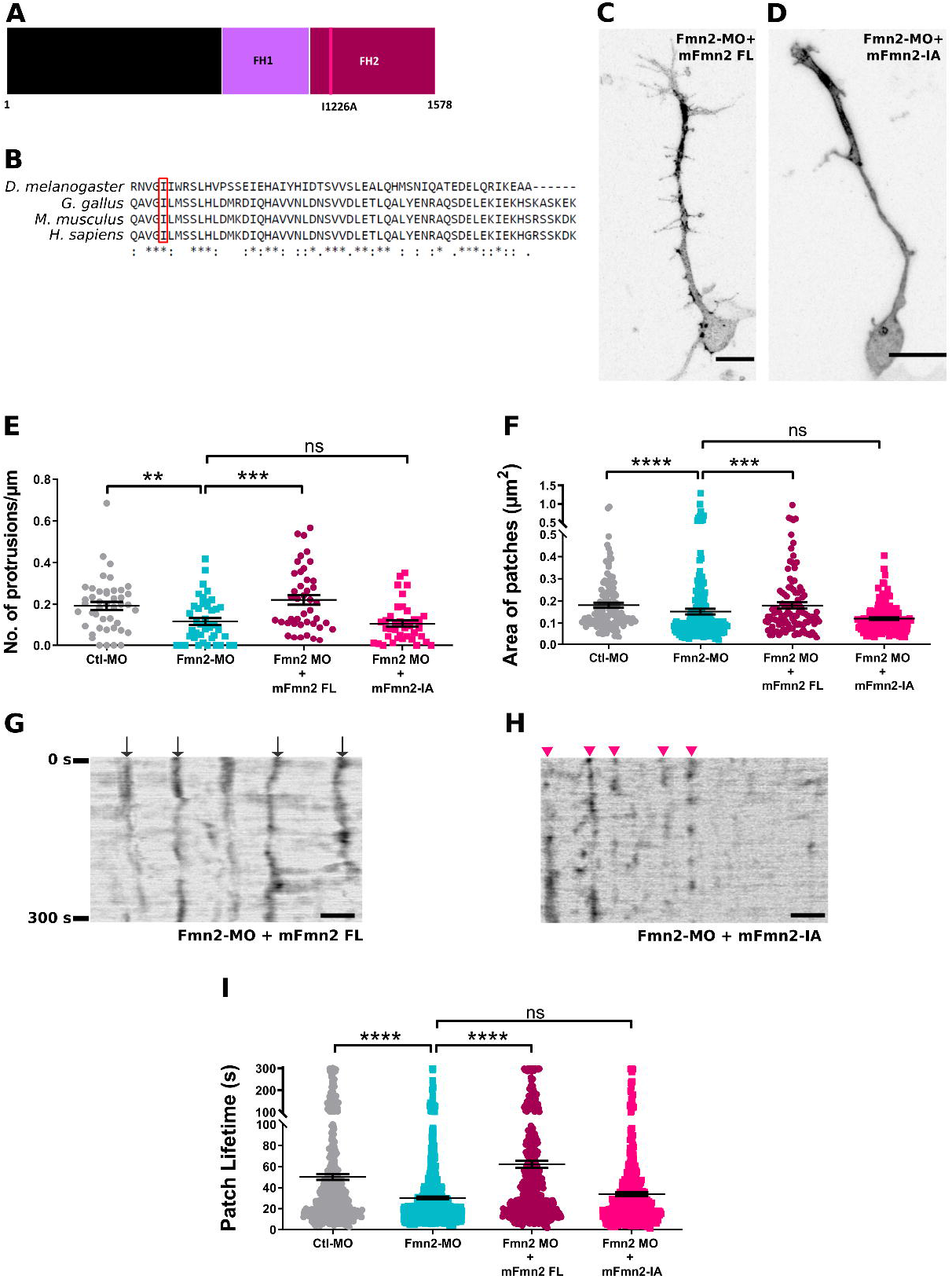
Actin nucleation activity of Fmn2 is necessary for the development of axonal protrusions and F-actin patch stability. (A) Schematic representation of domains of Fmn2 with the nucleation-dead mutation (I1226A) indicated in the FH2 domain .(B) Clustal-W multiple alignment of the FH2 domain of Fmn2 protein showing conservation of this critical amino acid (boxed) across phyla. (C-E) Representative micrographs of neurons with collateral protrusions co-expressing Fmn2-MO and GFP-tagged constructs of mouse full-length (mFmn2 FL) or the nucleation-dead mutant (mFmn2-I2A). (E) Quantification of protrusion density in Ctl-MO (n = 44), Fmn2-MO (n = 42, p=0.0032), Fmn2-MO + mFmn2-FL rescue (n = 42, p=0.0005), and Fmn2-MO + mFmn2-I2A rescue (n= 39, p=0.7485) transfected neurons. (F) The F-actin patch areas were rescued by morpholino-resistant full-length mouse Fmn2 (mFmn2 FL) but not the morpholino-resistant nucleation-dead mutant (mFmn2 IA). Ctl-MO n = 128; Fmn2-MO n = 189, p <0.0001; Fmn2-MO + mFmn2 FL n =110, p= 0.0001; Fmn2-MO + mFmn2-I2A n= 156, p=0.7276. (G,H) Representative kymographs indicating the lifespan of F-actin patches in Fmn2-MO+mFmn2 FL (G) and Fmn2-MO+mFmn2-I2A (H) transfected neurons. The pink arrow heads mark the ‘blinking’ phenomena of F-actin patches. (I) The quantification and comparison of patch lifetime in Ctl-MO (n = 490), Fmn2-MO (n = 693, p <0.0001), Fmn2-MO + mFmn2 FL (n = 456, p <0.0001), and Fmn2-MO + mFmn2-I2A (n= 522, p= 0.6537) transfected neurons. The data are obtained from at least three biological replicates. All the values were plotted with the mean +/-SEM indicated. The data were compared using the Kruskal-Wallis test followed by uncorrected Dunn’s test. ns, p>0.05;; * *, p<0.01; * * *, p<0.001; * * * *, p<0.0001. Scale bar: 10 µm (C,D) and 2 µm (G,H).

Axonal branching was rescued with the expression of morpholino resistant full-length mFmn2 in morphant chick neurons (Figure 4 C,E). However, the nucleation deficient mutant of mouse Fmn2 (mFmn2-IA) failed to rescue the decrease in collateral branching (Figure 4 D,E). The reduction in F-actin patch area and lifetime in Fmn2 morphant axons were also rescued by coexpression of mFmn2-FL but the mFmn2-IA failed to rescue either of these F-actin patch parameters (Figure 4 F-I) with small, short-lived actin patches.

Taken together, the rescue experiments demonstrate that the nucleation activity of Fmn2 is required for regulating F-actin patch dynamics and consequently collateral branching.

### Fmn2 and ADF act antagonistically to regulate actin patch stability and axonal branching

The F-actin patch ‘blinking’ observed upon depletion of Fmn2 suggested that axonal F-actin patches are dynamically balanced by stabilizing and destabilizing activities. With the identification of F-actin nucleation by Fmn2 in regulating F-actin patches, we sought to test this hypothesis directly. Formin nucleated F-actin is resistant to the actin depolymerising activity of cofilin both *in vitro* and *in vivo* (Mizuno et al., 2018). Further, recent evidence implicates the actin depolymerising factor (ADF)/cofilin family in regulating axon branching (Tedeschi et al., 2019). We tested if Fmn2 maintains patch stability by antagonising the depolymerizing activity of ADF (the ADF/cofilin family member in chick with highest depolymerizing activity (Chen et al., 2004).

Coexpression of a phospho-mimetic, inactive ADF (S3E) along with Fmn2-MO was able to rescue the reduction in collateral protrusion observed in axons depleted of Fmn2 (Figure 5A). Conversely, co-expression of constitutively-active ADF (S3A) along with Fmn2-GFP suppressed the increased branching observed upon Fmn2 overexpression (Figure 5B).

**Figure 5.**
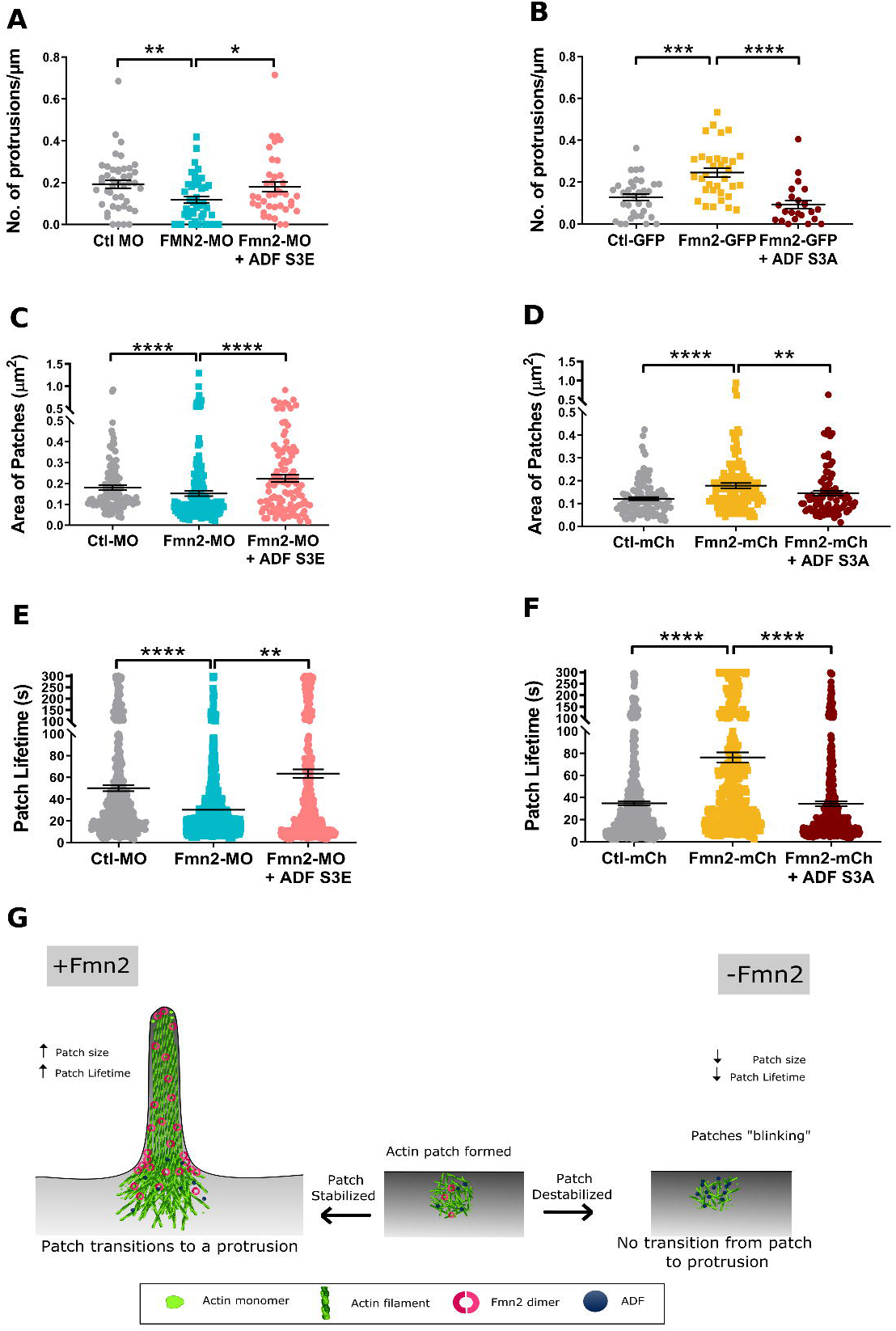
Antagonistic activities of Fmn2 and ADF dynamically regulate axonal protrusions and F-actin patch stability. (A) Comparison of axonal protrusion density in Ctl-MO (n =44), Fmn2-MO (n =42, p=0.002) and ADF-S3E rescue (n = 38, p=0.0482) transfected neurons. (B) Axonal protrusion density in Ctl-GFP (n = 36), gFmn2-GFP (n = 33, p= 0.0001), and ADF-S3A + Fmn2 GFP (n = 23, p <0.0001). (C) Comparison of F-actin patch areas in Ctl-MO (n =128), Fmn2-MO (n= 189, p<0.0001), and Fm2-MO + ADF-S3E (n = 106, p<0.0001) treated axons. (D) F-actin patch areas in Ctl-mCh (n = 121), Fmn2-mCh (n = 125, p<0.0001) and Fmn2-mCh + ADF S3A (n = 86, p=0.0069) expressing neurons. (E) F-actin patch lifetime in Ctl-MO (n = 490), Fmn2-MO (n = 693, p<0.0001), and ADF-S3E + Fmn2-MO (n = 483, p=0.0019) transfected neurons. (J) Graphical representation of F-actin patch lifetimes in Ctl-mCh (n = 576), gFmn2-mCh (n = 379, p<0.0001)) and ADF-S3A + gFmn2 (n = 476; p<0.0001) expressing neurons. (G) Model of Fmn2 in F-actin patch dynamics leading to axonal collateral protrusion. An actin patch is initiated by nucleators such as Arp2/3 and/or other formins and is maintained by the activity of Fmn2, which insulates the patch from ADF-mediated depolymerisation. Stability of the actin patch renders it competent for signal-induced transition to a nidus of actin polymerisation and elongation that initiates a collateral protrusion. On the contrary, in the absence of Fmn2, the actin patch is is destabilized by actin severing factors such as ADF, leading to patch dissipation without any protrusion initiation. All values were plotted with the mean +/-SEM indicated and were obtained from at least three independent experiments. The Kruskal-Wallis test followed by uncorrected Dunn’s test for multiple comparisons was used to compare across treatments. *, p<0.05; * *, p<0.01; * * *, p<0.0001; * * * *, p<0.0001.

Consistent with the rescue of branching, expression of the inactive ADF S3E was also able to rescue the reduction patch area and lifetime in Fmn2 morphants (Figure 5 C,E). Similarly, the expression of constitutively-active ADF S3A restored the Fmn2 overexpression-induced increased actin patch area and lifetime (Figure 5 D,F) to control levels.

Thus, a dynamic balance between Fmn2-mediated F-actin assembly and ADF-induced disassembly regulates the lifetime of axonal F-actin patches and, in turn, the development of axon collateral branches.

## DISCUSSION

Co-existence of functionally distinct cytoskeleton networks in the cytosol necessitates unique regulatory mechanisms that confer specific properties to these networks. The F-actin structures in the axon shaft include the membrane-associated periodic skeleton (MPS), vesicle-associated F-actin ‘hotspots’ and ‘trails’ and collateral branch-inducing focal F-actin patches (Leterrier et al., 2017; Mutalik and Ghose, 2020). The initiation of neurite collateral branching by filopodia-like protrusions generated by F-actin filaments emanating from a focal accumulation of F-actin in the neurite shaft (F-actin patch) is well established in both axons (Armijo-Weingart and Gallo, 2017; Gallo, 2006; Ketschek and Gallo, 2010; Orlova et al., 2007) and dendrites (Nithianandam and Chien, 2018; Sturner et al., 2019). However, the regulation of F-actin patch dynamics remains poorly understood.

Studies in vertebrate sensory neuron axons and fly dendrites have identified WASP-dependent induction of F-actin nucleation by Arp2/3 in initiating the formation of F-actin patches (Spillane et al., 2011; Sturner et al., 2019). The formins are speculated to be involved in filament elongation, though experimental evidence of their involvement in axonal branching is lacking. In this study, we demonstrate that the formin, Fmn2, is a key regulator of F-actin patch dynamics. Fmn2 antagonises the F-actin depolymerising activity of ADF to maintain F-actin patches and consequently regulates the initiation of axon collateral branching.

We find that Fmn2 expression is positively correlated with axonal protrusions, and Fmn2 localises to protrusion-initiating F-actin patches. However, depletion or overexpression of Fmn2 does not change the number of F-actin patches in axons. This observation indicates F-actin patch formation to be independent of Fmn2 activity. Fmn2 knockdown results in short-lived patches that iteratively disappear, recover and then diminish again at the same location within the axon. This ‘blinking’ phenomenon suggests a role for Fmn2 in F-actin patch maturation or maintenance. It appears that a nucleating core is established independent of Fmn2, however, the maturation and stability of this F-actin network is Fmn2 dependent. This hypothesis is consistent with our observation that F-actin nucleation activity of Fmn2 is necessary for its function in F-actin patch dynamics. Overexpression of Fmn2 was found to increase F-actin patch lifetimes, underscoring Fmn2 function in patch stability.

Arp2/3 activity-dependent F-actin patch initiation has been described in multiple systems, including chick sensory neurons, and consistently these networks have a dendritic architecture (Spillane et al., 2011). Thus, it appears Arp2/3, perhaps working in tandem with other actin nucleation promoting factors, is critical for patch formation. However, other activities, including that of Fmn2, is required to aid patch elaboration and maintenance.

Longer-lasting patches are likely to have an increased probability of initiating actin structures required to generate membrane protrusions (Loudon et al., 2006). Comparison of F-actin patch sizes and lifetimes support this hypothesis with patches converting to protrusions being larger and longer-lasting. In fact, the few protrusion that form in axons depleted of Fmn2 also originate from the few remaining patches with longer lifetimes.

The ADF/cofilin family of F-actin severing and disassembly factors are essential for actin turnover and involved in the dynamics of diverse F-actin networks(Bamburg and Bernstein, 2010; Bernstein and Bamburg, 2010; Kanellos and Frame, 2016). Yeast ADF/Cofilin is a critical component of cortical F-actin patches (Chen and Pollard, 2013; Lappalainen and Drubin, 1997) and recently, loss of ADF/Cofilin activity has been shown to increase axonal branching (Tedeschi et al., 2019). These observations prompted us to directly test if Fmn2 antagonises ADF/Cofilin function.

We expressed dominant-negative and constitutively activated mutants of chick ADF - the major ADF/cofilin family member in the chick nervous system (Chen et al., 2004; Devineni et al., 1999) – while manipulating Fmn2 expression. Inhibition of ADF activity in the background of Fmn2 depletion restored the reduction in F-actin patch stability and rescued the deficit in initiating axonal protrusions. Conversely, Fmn2-overexpression induced increased stability of F-actin patches and protrusive activity were both reduced to control levels upon co-expression of constitutively-activated ADF. These results strongly implicate antagonistic activities of ADF and Fmn2 in regulating F-actin patch dynamics and, in turn, the initiation of axonal protrusions (Figure 5G). While the exact molecular mechanism of this antagonism remains unknown, recent studies have demonstrated that formin-generated F-actin is resistant to ADF/cofilin-mediate severing both *in vitro and in vivo* (Mizuno et al., 2018). As ADF/cofilin activity is ubiquitous in eukaryotic cells, dynamic antagonism by F-actin assembly activities, like that of Fmn2 shown here, is likely to be widespread across phyla in a range of cellular processes. The actin nucleator Spire, known to function cooperatively with Fmn2 (Montaville et al., 2014; Vizcarra et al., 2011), has been shown to regulate branching in dendritic arbours (Ferreira et al., 2014) though the underlying molecular mechanism has not been elucidated. Taken together with our observations, it is possible that Fmn2 and Spire collaborate to regulate neurite branching in vertebrates.

We propose a dynamic balance of activities, in part involving Fmn2 and ADF, in the maintenance and dynamics of protrusion-competent axonal actin patches (Figure 5G). Resistance to cofilin-induced severing observed in formin-nucleated actin filaments (Mizuno et al., 2018) may have a role in mixed F-actin networks co-regulated by both Arp2/3 and formins, like the axonal F-actin patches. Recent observations from Arp2/3 and formin-induced mixed networks suggest cooperativity where formin-induced filament elongation protects the dense Arp2/3-induced dendritic networks from severing activity of cofilin (Bleicher et al., 2020). In addition to the Fmn2-ADF antagonism described here, Fmn2 may regulated F-actin elongation by competing with actin filament capping activities.

Other functionally distinct F-actin networks display different modes of regulation. In the proximal end of the axon, specialized F-actin patches form a selective cargo filter for axonal proteins. However, formin activity is dispensable for these structures and neither does Fmn2 localise to these proximal axon networks in the (Balasanyan et al., 2017). Similarly, Arp2/3-assembled cortical actin patches in yeast are insulated from formins by the activity of capping proteins to maintain a distinct identity (Billault-Chaumartin and Martin, 2019).

In summary, we demonstrate that the neurodevelopmental disorder-associated formin, Fmn2, regulates the maintenance and stability of a specific cytoskeleton network that is critical for the initiation of axon collateral branching. We identify an antagonistic mechanism between Fmn2-mediated F-actin assembly and disassembly by ADF in regulating the dynamics of axonal F-actin patches that are essential for axonal branching.

## MATERIALS AND METHODS

### Culturing primary neurons

Spinal cord tissue from stage HH 25-26 (Day 5-6) chick embryos (Venkateshwara hatcheries Ltd.) were dissected out in L-15 medium (Gibco, HiMedia) with added 1x penstrep (Gibco). After dissection, the tissue was trypsinized (0.05% Trypsin-EDTA (Lonza, HiMedia) at 37°C for 15-20 minutes and then resuspended in Optimem® (Gibco) prior to cuvette-based electroporation (CUY21SC, Nepagene). The transfected neurons were plated on glass-bottomed 35 mm plastic dishes and incubated at 37°C for 24-36 hours in L-15 medium with 10% fetal bovine serum (FBS; Gibco) and 1x penstrep (Gibco). The dishes were coated with poly-L-lysine (1 mg/ml, Sigma) and laminin (20 µg/ml) (Sigma) before plating.

### Transfection with plasmids and morpholinos

The details of the plasmids used in this study are given in Table S1. Fmn2 was knocked down in cultured neurons using translation-blocking anti-gFmn2 morpholinos (Fmn2-MO; 5’ CCATCTTGATTCCCCATGATTTTTC 3’) (100 μM). Standard control morpholino (Ctl-MO; 5’ CCTCTTACCTCAGTTACAATTTATA 3’) was used as a negative control. These reagents have been extensively characterised for knockdown efficiency and specificity (Ghate et al., 2020; Sahasrabudhe et al., 2016). For fixed experiments, the neurons were co-transfected with pCAG-GFP and Ctl-MO or Fmn2-MO. Correspondingly, for live-imaging experiments, pCAG-tractin-GFP was used in place of pCAG-GFP. Rescue experiments were performed with triple transfections of pCAG-GFP + Fmn2-MO + mCherry-tagged constructs.

For fixed analysis of Fmn2 overexpression experiments, pCAG-gFmn2-GFP was co-transfected with either pCAG-mCherry or with pCAG-ADF S3A-mCherry. For live-imaging of Fmn2 overexpression, pCAG-tractin-GFP was co-transfected with either pCAG-mCherry or gFmn2-mCherry or gFmn2-mCherry+ ADF S3A mCherry.

### Immunofluorescence

For quantifying branching, the neurons were fixed in 4% paraformaldehyde, 0.25% glutaraldehyde in PHEM buffer (60 mM PIPES (Sigma), 25 mM HEPES (Sigma), 10 mM EGTA (Sigma), and 4 mM MgSO4·7H20). After 10 minutes of permeabilization in 0.2% triton-PHEM, blocking was done with 3% BSA for 1 hour at room temperature. Phalloidin-633 (Invitrogen) was used in 1:250 concentration to label F-actin before mounting in MOWIOL-DABCO (2.5% 1, 4-Diazabicyclo-octane (DABCO) (Sigma), 10% MOWIOL 4-88 (Sigma), 25% glycerol (Sigma) and 0.1M Tris-HCl, pH-8.5).

### Imaging

The fixed images were acquired using Zeiss LSM 710, 780 and Leica SP8 confocal systems with 63x, 1.4 NA oil objective and in every system.

Live-imaging of actin patches was done in widefield mode on the Olympus IX-81 system (Olympus Corporation, Tokyo, Japan) with 100x, 1.4 NA Apo oil immersion objective and Hamamatsu ORCA-R2 CCD camera. The temperature was maintained at 37°C in a humid chamber for the duration of imaging. Time-lapse images of Tractin-GFP were acquired using the RT Xcellence software with a time interval of 2 s for 150 frames.

For colocalisation of Fmn2 and actin, imaging was done using the Oxford Nanoimager system with HILO (High Inclination and Laminated Optical sheet) setup using the 100X NA objective. Dual-colour images were acquired with 500ms time-interval for 600 frames at 37 °C.

### Image analysis

For quantification of protrusion density, the entire axon was considered, excluding the growth cone and the soma. Protrusions ≥ 0.5µm were considered for analysis. The first frame of each tractin-GFP time-lapse movie was utilized to calculate the patch area and density. The area of a F-actin patch was measured manually using the freehand selection tool after background subtraction. For calculating patch area of patches that formed protrusions, the area of the patch one frame before a detectable protrusion was observed, was taken into consideration. For patch lifetime analysis, the time-lapse images were drift corrected using the StackReg plugin of ImageJ followed by generation of kymographs using the KymoClear 2.0 plugin. Segmented lines were drawn manually on each kymograph to mark the patches. Patch lifetime was calculated using the velocity-measurement tool plugin. For calculating the lifetime of patches that gave rise to a protrusion, the patches were tracked manually from the first frame of detectable appearance to the last frame of disappearance. For colocalisation analysis, ROI’s of actin patches were identified using the tractin channel. Then, fluorescence of every ROI in both the channels was quantified using the formula, Corrected fluorescence (CTCF) = Integrated Density – (Area of selected cell X Mean fluorescence of background readings). The fluorescence values of both the channels were plotted in X-Y table in Prism 8 to calculate Pearson’s correlation coefficient and r-squared values. All analysis was performed after anonymizing the genotype.

### Statistical analysis and graphical representation

All the graphical representation and statistical analyses were performed using GraphPad Prism 8. Graphical representations were plotted using scatter dot plot with middle line showing the mean and error bars indicating the standard error of mean (SEM). Mann-Whitney U test was employed for two groups and for multiple comparisons Kruskal-Wallis test was done followed by uncorrected Dunn’s test as indicated in each figure. Number of data points quantified in each graph and the biological replicates have been indicated in the figure legends.

## ETHICS APPROVAL

All protocols used in this study were approved by Institutional Animal Ethics Committee and the Institutional Biosafety Committee of IISER Pune.

## AVAILABILITY OF DATA AND MATERIAL

All data generated or analysed during this study are included in this published article. The raw data are available from the corresponding author on reasonable request.

## AUTHORS’ CONTRIBUTIONS

Conceptualization: T.K. and A.G.; Investigation and analysis: T.K., S.S.D., D.S.K. and L.K.S.; Writing of manuscript: T.K. and A.G.; Funding Acquisition: A.G.. All authors gave final approval for publication and agree to be held accountable for the work performed therein.

## ACKNOWLEDGMENTS

The authors thank Dr. N. K. Subhedar, IISER Pune for his critical reading of this manuscript and Dhriti Nagar for assistance in making the graphics for the model. The authors acknowledge the IISER Pune Microscopy Facility and the National Facility for Gene Function in Health and Disease (NFGFHD) at IISER Pune for access to equipment and infrastructure.

