## Supplementary information for "Antagonistic activities of Fmn2 and ADF regulate axonal F-actin patch dynamics and the initiation of collateral branching"

**This file includes:**

Figure S1
Table S1

**A**

| Treatment | Area of total no. of patches (µm²) | Area of patches that form protrusion (µm²) |
|---|---|---|
| Ctl-MO | 0.1805± 0.012 | 0.2749± 0.038 |
| Fmn2-MO | 0.1519± 0.012 | 0.3073± 0.039 |

**B**

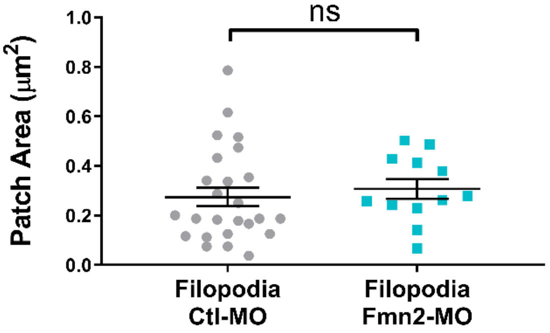


**C**

| Treatment | Lifetime of total no. of patches (s) | Lifetime of patches that form protrusion (s) |
|---|---|---|
| Ctl-MO | 50.10 ± 2.84 | 60.35 ± 10.71 |
| Fmn2-MO | 30.16 ± 1.29 | 64.13 ± 14.71 |

**D**

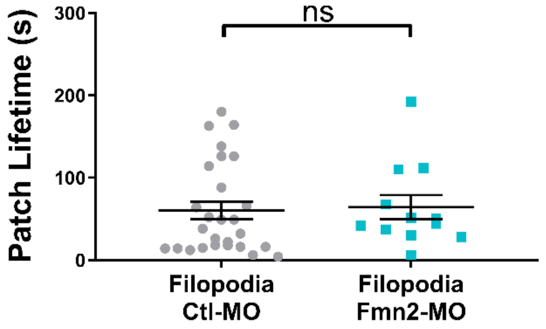


**Figure S1. Patches giving rise to filopodia have larger areas and higher lifetimes.**

(A and C) Tables showing the mean patch area (B) and patch lifetime (D) of all the patches and patches that form filopodia in both Ctl-MO and Fmn2-MO transfected axons. (B and D) Quantification of patch dynamics in Ctl-MO and Fmn2-MO treated neurons. Comparison of patch area (A) and patch lifetime (C) that form filopodia in both Ctl-MO (n = 25 and 27, respectively) and Fmn2-MO (n = 12; p-value = 0.3392 and 0.5036, respectively; ns p>0.05) treated neurons. All the values are plotted as mean and SEM and data were analyzed using Mann-Whitney U test and obtained from at least three independent experiments

**Table S1: Plasmid constructs used in this study indicating source, vector backbone, cloning strategy and primers used for cloning.**

| Plasmid Name | Plasmid Backbone | Insert Name | Source | Gene bank acc. no. | Primer used for cloning | Region cloned and cloning strategy |
|---|---|---|---|---|---|---|
| pCAG-GFP | pCAG | - | Addgene (11500) | - | - | - |
| pCAG-mCherry | pCAG | mCherry | This study | - | Fwd 5'ATATATACCGGTCGCCACCATGGT GAGCAAGGGCGAGGAGG3' Rev 5'ATATATGCGGCCGCTTTACTTGTAC AGCTCGTCCATGCCGCC3' | cloned by ligation between AgeI and NotI sites |
| pCAG-Tractin-GFP/mCherry | pCAG-GFP and pCAG-mCherry | F-tractin | This study | NM_031045.2 | Fwd 5'ATGGCGCGACCACGGGGCGCGGGGCCCTGCAGC CCCGGGTTGGAGCGGGCTCCGCGCCGGAGCGTCG GGGAGCTG3' Rev 5'CCCCCTGCGGCCGCTGCGGCGGCGACTGCGGCG CAGCGCGCTTCGAAGAGCAGGCGCAGCTCCCCGAC GCTCCGG3' | 1-113, Cloned by homologous recombination for between Age1 and Kpn1 |
| pCAG-gFmn2-GFP/mCherry | pCAG-GFP and pCAG-mCherry | gFmn2 | Jacob et al., 2016 | KU711529.1 | NA | complete cds, cloned by Homologous recombination using SmaI |
| pCAG-mFmn2-FL-GFP/mCherry | pCAG-GFP and pCAG-mCherry | mFmn2; Gift from Dr. Philip Leder, Harvard Medical School | This study | NM_019445 | Fwd 5'AGTCGACGGTACCCGCCACCATGG GGAACCAGGATGGGAAG 3' Rev 5'GGTGGCGACCGGTGGATCCCGGGt CGTTTTCATGCTTATCTTCGCTTTAAT C 3' | complete cds, cloned By homolous recombination between KpnI and XmaI |

| | | | | | | |
|---|---|---|---|---|---|---|
| pCAG-mFmn2-ΔFSI-GFP/mCherry | pCAG-GFP and pCAG-mCherry | pCAGmFmn 2-FL | This study | NM_019445 | Fwd 5'GCAAAGAATTCTGCAGTCGACGGT ACCCGCCACCATGGGGAACCAGGAT GG 3' Rev 5'CCATGGTGGCGACCGGTGGATCCC GGGCCTTCTGCCTACACACCTCC3' | 1-4662, cloned by Homologous recombination between KpnI and XmaI |
| pCAG-mFmn2-I2A-GFP/mCherry | pCAG-GFP and pCAG-mCherry | pCAGmFmn 2-FL | This study | NM_019445 | Fwd 5'CAAAAGGTCACAAGCAGTAGGAGCTCTAATGTCTA GTCTGCATTTAGATATG3' Rev 5'CATATCTAAATGCAGACTAGACATTAGAGCTCCTA CTGCTTGTGACCTTTTG3' | Mutation I1226A, cloned by homologous recombination between KpnI and XmaI |
| pCAG-ADF S3E-GFP/mCherry | pCAG-GFP and pCAG-mCherry | ADF S3E, Gifted by Dr. J. Bamburg, Colorado State University, USA | This study | - | Fwd 5'ATTTTGGCAAAGAATTCCACCGCCATGGCAGAAGG AGTACAAG3' Rev 5'CTTTTGAAGGAAGTCCTGTGgCCCGGGATCCACCG GTCGC3' | complete cds, cloned by homologous recombination between Age1 and Kpn1 |
| pCAG-ADF S3A-GFP/mCherry | pCAG-GFP and pCAG-mCherry | Gifted by Dr. J. Bamburg, Colorado State University, USA | This study | - | Fwd 5'ATTCTGCAGTCGACGGTAC3' Rev TCACCATGGTGGCGACCGGTGGATCCCCCACAGGA CTTCCTTCAAAAGC3' | complete cds, cloned by homologous recombination between Age1 and Kpn1 |
